# Motor Learning Driven by Sensory Prediction Errors is Insensitive to Task Performance Feedback

**DOI:** 10.1101/2025.06.28.662119

**Authors:** Gaurav Panthi, Pratik Mutha

## Abstract

Accurate motor behavior relies on our ability to refine movements based on errors. While sensory prediction errors (SPEs), mismatches between expected and actual sensory feedback, predominantly drive such adaptation, task performance errors (TPEs), or failures in achieving movement goals, also appear to contribute. However, whether and how TPEs interact with SPEs to shape net learning, remains controversial. This controversy stems from difficulties in experimentally decorrelating these errors, ambiguity related to possible interpretations of task instructions, and inconsistencies between theory and computational models. To try and resolve this issue, we employed variants of an “error-clamp” adaptation paradigm across four reaching experiments (N = 144). Addressing the ambiguity of whether or not the TPE is indeed ignored in standard error-clamp designs as assumed in theoretical (but not computational) models, Experiment 1 explicitly manipulated TPE magnitude by shifting the endpoint feedback location while holding SPE constant. We found that learning was uninfluenced by TPE size. Experiment 2 assumed that the TPE is in fact disregarded under clamp instructions. To then study SPE-TPE interactions, we induced TPEs of varying magnitudes by shifting the target location (“target jump”) while always clamping feedback to the original target location. Here, instructions to reach the new target also induced an SPE. Crucially, learning driven by this SPE was again unaffected by TPE magnitude, a result validated by two additional experiments. Our findings consistently demonstrate that SPE-mediated learning remains impervious to variations in task performance feedback, and point to a distinction in learning mechanisms triggered by these two error signals.

## INTRODUCTION

The constantly changing environment we interact with challenges our sensorimotor system in different ways. For instance, environmental perturbations introduce errors in our actions, which, if left uncompensated, can result in a failure to achieve intended movement goals. The sensorimotor system demonstrates an ability to quickly recalibrate motor plans to compensate for such errors if they occur repeatedly, a process termed motor adaptation. There has been growing interest in motor neuroscience in understanding the kind of errors that environmental perturbations induce and the compensatory learning or adaptation mechanisms they stimulate for such recalibration to occur. Laboratory experiments of motor adaptation employing visual (Kumar et al., 2020; Martin et al., 1996; Morehead et al., 2017) or dynamic (Sainburg et al., 1999; Shadmehr & Mussa-Ivaldi, 1994) perturbations have focused largely on two types of error signals: a sensory prediction error (SPE) and a task performance error (TPE). An SPE is an execution related error, operationalized as a mismatch between actual and predicted sensory feedback. On the other hand, TPEs are related to task success and defined as a failure in achieving the task goal, such as a feedback cursor failing to strike a reach target. Our main goal in the current work is to better understand the interactions between the TPE and the SPE during motor adaptation, an issue of tremendous research interest in recent times, but often laden with inconsistent and controversial outcomes (Al-Fawakhiri et al., 2023; Kim et al., 2019; Leow et al., 2018; Oza et al., 2024; Tsay et al., 2022).

A fundamental problem in using canonical adaptation paradigms for addressing this issue is that in such tasks these two errors end up being inherently correlated, posing a challenge in isolating their individual contributions or studying their interaction in a meaningful way. For example, in a visuomotor rotation adaptation paradigm, in which the visual feedback of the movement (often a cursor displayed on a screen) is rotated relative to actual hand motion, the magnitude of the TPE is tightly linked to that of the SPE, and a reduction in the SPE through learning leads to a concomitant decrease in the TPE. This limitation can be mitigated to some degree by employing “error clamps”, in which an SPE is induced by rotating the visual feedback of hand motion (cursor), but de-yoking cursor motion from the actual hand movement. In these tasks, participants are made to understand that they should ignore this error and move their hand to the target, yet they exhibit gradual drift of the hand away from the target and robust aftereffects (Morehead et al., 2017). However, the error-clamp task also typically engenders an inherent TPE since the cursor consistently misses the target. Crucially, whether participants also ignore this TPE when given the instruction to ignore the cursor remains unknown, particularly for two reasons: 1) the amount of learning in response to the same clamped SPE appears to be greater when the feedback cursor misses the target (TPE present) compared to when it does not (TPE absent) (Al-Fawakhiri et al., 2023; Leow et al., 2018; Oza et al., 2024; Tsay et al., 2022) and 2), an explicit TPE term is included when computationally modeling the behavior exhibited by participants in the error-clamp paradigms (Kim et al., 2019; Tsay et al., 2022), acknowledging its presence. Yet, learning in error-clamp tasks is said to be driven entirely by the SPE, creating a discrepancy between theory and modeling. Hence, the suitability of the typical clamp paradigm for studying interactions between TPE and SPE during learning, which is our primary intent here, remains an open question.

To better address this issue, we begin by considering two possibilities. First, we consider the possibility that in a typical error-clamp paradigm, the TPE is unambiguously present and not ignored. If this is the case, then the standard clamp paradigm is not ideal for probing the interactions between SPE- and TPE-mediated changes since these two error signals remain not only correlated but also unchanged as the task progresses. To overcome this limitation, in our first set of experiments, we modified the error-clamp paradigm such that the clamped cursor is visible only until the participant moves halfway to the reach target and then disappears. This enables the participants to perceive the induced SPE. In addition, we vary the magnitude of the TPE by re-displaying the cursor at the reach endpoint, but at different distances relative to the target location. Thus, by keeping SPE fixed and systematically varying the TPE, this new paradigm enables us to investigate if and how TPEs (of different magnitudes) influence SPE-mediated learning. To reiterate, the critical assumption here is that when given instructions to ignore the cursor as in the standard clamp design, the TPE is *not* ignored and *can* influence the change in hand angle brought about by the SPE.

The second possibility that we consider is that the TPE is indeed ignored when participants are instructed to disregard the cursor feedback in the clamp paradigm. Now, if the TPE is made irrelevant by asking participants to ignore it, and they indeed do so, then again, the standard clamp paradigm becomes inadequate to probe SPE-TPE interactions. We therefore turned to a new paradigm we recently developed wherein we shift the location of the reach target (target jump) while clamping the feedback cursor in the direction of the original target (Oza et al., 2024). Since the cursor always misses the “jumped” target, a conspicuous TPE is generated. Furthermore, in this task, by instructing participants to aim to the new target location while clamping the motion of the cursor towards the old one, an SPE can be induced “on the fly”. To bring forth the influence that TPEs might have on the learning that this SPE induces, one could then manipulate the amplitude of the target jumps and instruct participants to respond to them by aiming to the new target location. We pursue this goal in our second set of experiments. Note that Tsay and colleagues (Tsay et al., 2022) have also used a similar paradigm to probe this issue. However, their paradigm only allowed the measurement of trial-by-trial changes since the average error is kept zero. In contrast, our approach enables us to investigate the effect of the TPE on the net adaptive response that emerges with time and practice.

In sum, we employ two distinct yet complementary approaches to study the interaction between TPE and SPE. Across our experiments, our results indicate that SPE-mediated learning remains robustly uninfluenced by variations in TPE, strengthening the idea of impervious nature of SPE and the learning mechanism it engages.

## MATERIALS AND METHODS

### Participants

A total of 144 young, healthy, right-handed adults (age range: 18-35 years) were recruited for different experiments in this study. Handedness was assessed via the Edinburgh Handedness Inventory (Oldfield, 1971). All participants provided written informed consent to participate in the study and were monetarily compensated for their time. The experimental protocols were approved by the Institute Ethics Committee of the Indian Institute of Technology Gandhinagar.

### Experimental setup

The experiments in this study were conducted using two different setups. For our first experiment, we used the Kinarm end-point robotic manipulandum (BKin Technologies, Canada) which consisted of a virtual reality system with a high-definition TV screen mounted horizontally above it (Figure 1A). Participants sat in an adjustable chair, facing the Kinarm system, and held the handle of a robotic arm, which allowed planar movements only. A mirror positioned between the robot and the TV display reflected the task presented on the screen while obstructing the direct vision of the hand. A circular cursor displayed on the screen provided visual feedback regarding the position of the hand (handle of the robotic system). This setup also allowed us to manipulate and differentiate the motion of the displayed cursor from the actual hand movement. Our remaining experiments were conducted using a digitizing tablet system (GTCO Calcomp, Scottsdale, AZ), which also employed a pseudo virtual reality setup similar to the Kinarm but differed in how movements were recorded (Figure 2A). Here, participants made planar movements on the tablet surface using a hand-held stylus, with a cursor on the screen providing visual feedback of their hand position. Again, a mirror placed between the tablet and the screen blocked vision of the arm. This too enabled us to dissociate the displayed location of the hand (cursor) from its actual position.

**Figure 1:**
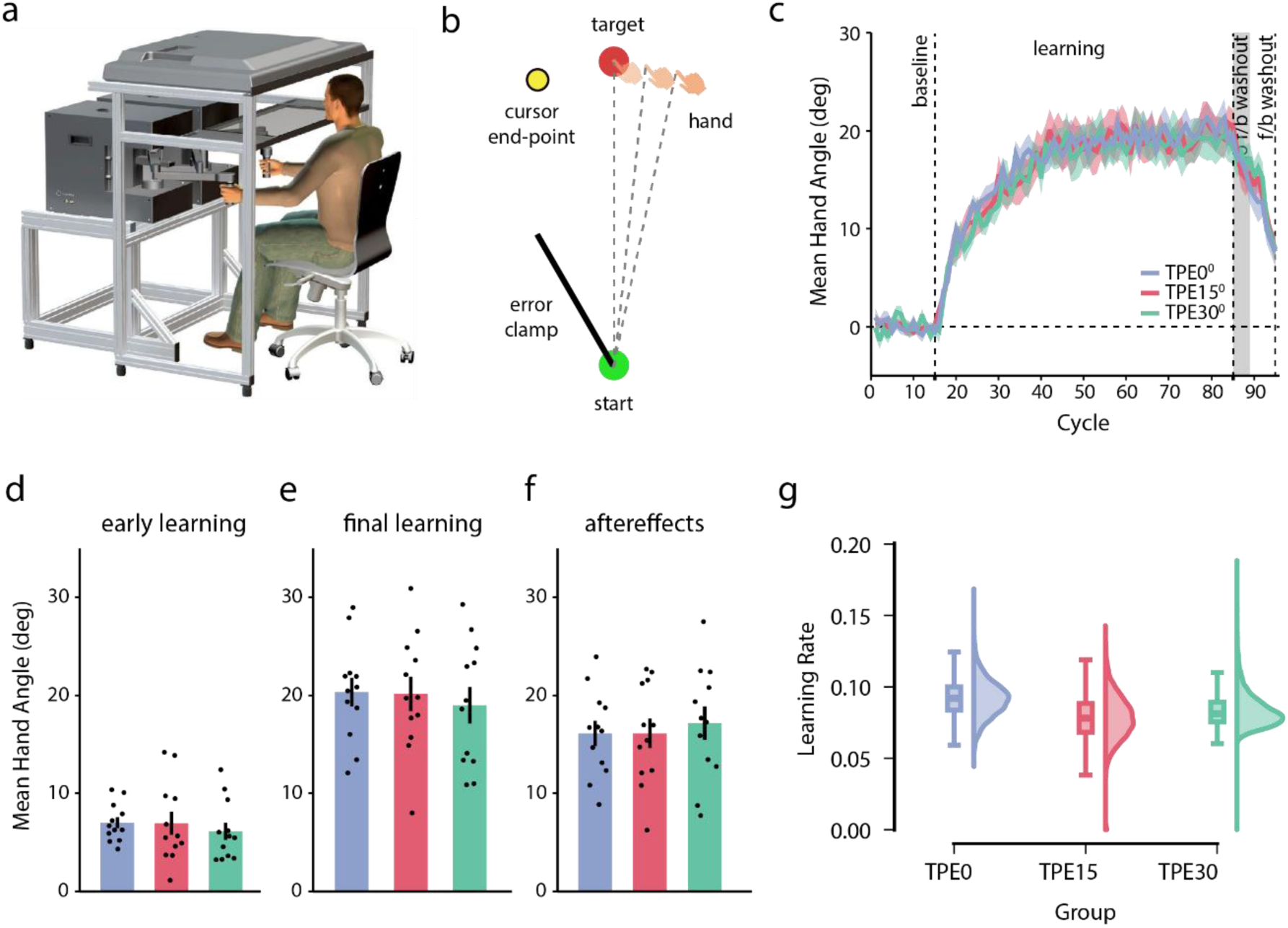
Experiment 1. **(a)** Participants performed reaching movements while holding the handle of a robotic manipulandum (Kinarm End-point). The start position, reach targets, and cursor feedback were displayed on a screen and reflected onto a mirror placed horizontally between the screen and the hand, occluding direct vision of the arm. **(b)** Task: Participants reached from the start position to the target. Feedback was provided by means of a cursor (yellow circle), but motion of the cursor was clamped at a 30° offset from the target direction. Additionally, the cursor was extinguished once the hand crossed a 6 cm distance, and was re-displayed upon movement completion as an endpoint. The endpoint cursor location could be on the target, or offset by 15° or 30° from the target, thereby creating task performance errors of different sizes. **(c)** Group-averaged hand deviations across cycles for the three experimental groups. Shaded regions represent SEM. **(d–f)** Mean ± SEM hand deviation during (d) early learning, (e) final learning, and (f) no-feedback washout phases. Dots represent individual participants. **(g)** Distributions and box-plots of learning rates for the three groups derived from exponential fits to 10000 bootstrapped hand deviation data samples.

**Figure 2:**
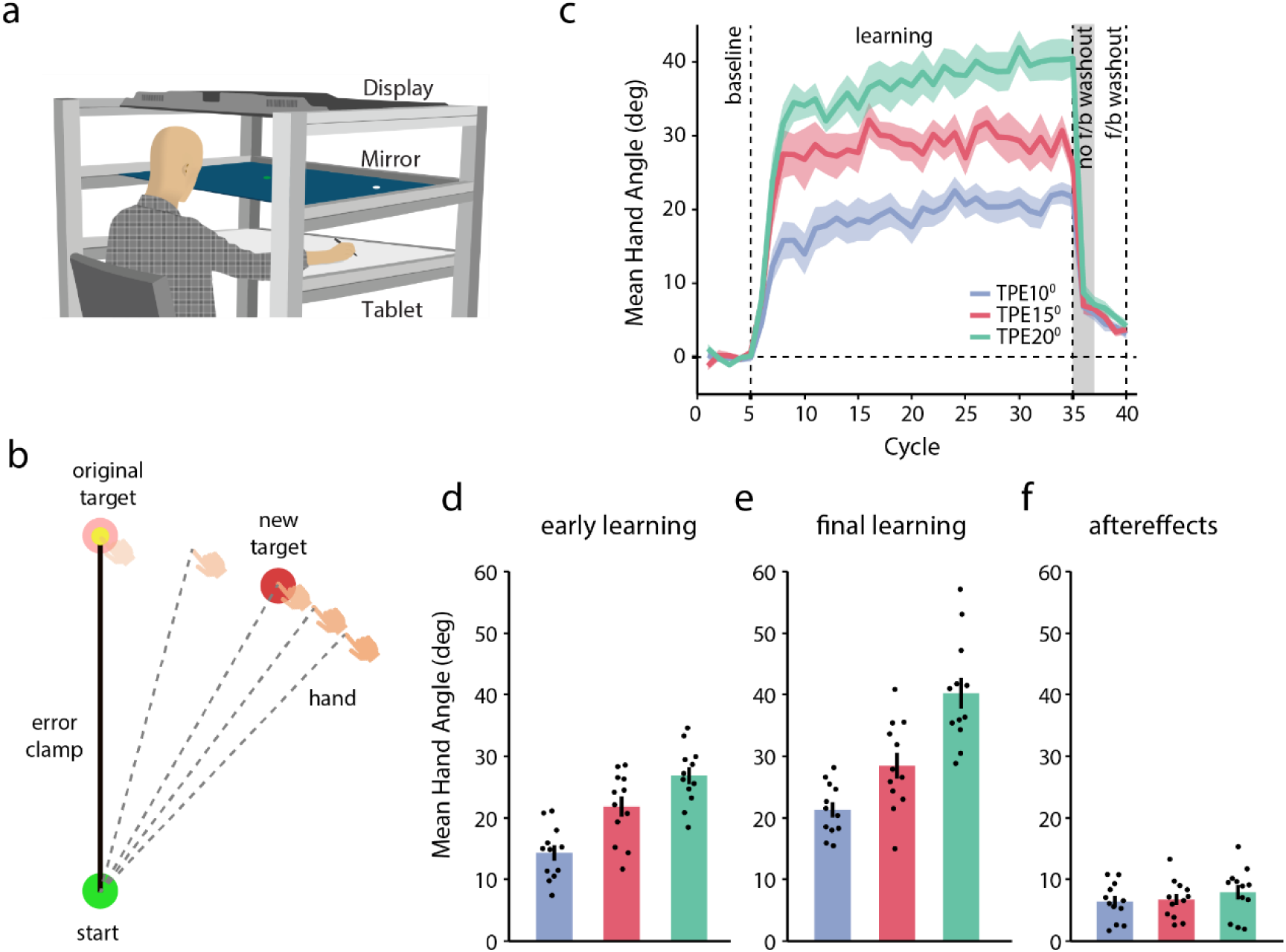
Experiment 2. **(a)** Setup for Experiments 2–4. Participants performed reaching movements on a digitizing tablet using a handheld stylus. The start position, reach targets and cursor feedback were displayed on a screen and reflected on a horizontal mirror placed above the tablet. **(b)** Task: Upon receiving the go cue, participants began a reach, but the originally displayed target was “jumped” from its original location to a new location either 10° (group 1), 15° (group 2), or 20° (group 3) away, creating task performance errors (TPEs). At the same time, the cursor was kept clamped in the direction of the original target. Since participants were instructed to reach to the new target, the mismatch between the expected motion of the cursor (aligned with the hand) and the actual motion of the cursor (clamped towards the original target) induced an SPE. **(c)** Group-averaged hand deviation across cycles for the three groups. Shaded regions represent SEM. **(d–f)** Mean ± SEM hand deviation during (d) early learning, (e) final learning, and (f) no-feedback washout phases. Dots represent individual participants.

### Task Design

All our experiments required participants to make point-to-point reaches from a stationary start circle to a target with their dominant, right arm. To begin a trial, participants first brought the cursor inside the start circle. After a delay of 500 milliseconds, a circular target appeared on the screen and participants made a quick reach towards it. During these movements, participants received one of three types of cursor feedback: veridical, clamped or no feedback at all. On veridical feedback trials, the cursor accurately represented the hand position, whereas the cursor remained invisible during no-feedback trials. On error-clamp trials, the cursor followed an invariant trajectory rotated by a specific angle, regardless of the direction in which the participant moved their hand. Importantly, participants were explicitly informed about the nature of the cursor feedback on these trials and were instructed to ignore it, focusing instead on moving their hand toward the target.

#### Experiment 1

In experiment 1, participants (n = 36) performed 380 reaches originating from the start circle (2 cm diameter) to one of four targets (2 cm diameter) located at a distance of 12 cm at one of the locations (45°, 135°, 225° and 315°) arranged around an imaginary circle. Target presentation was pseudorandom with a target appearing once in a cycle of 4 trials. The task consisted of 3 continuous blocks of trials: baseline (60 trials), learning (280 trials) and washout (40 trials). The experiment began with a block of familiarization trials to get participants to understand the task better. Following familiarization, participants performed baseline trials, comprised of 3 sub-blocks of 20 trials each: no feedback, partial feedback, and veridical feedback. In the “no feedback” sub-block, the cursor was not shown during the reach. In the “partial feedback” sub-block, the cursor remained visible only up to half the distance to the target (6 cm) and then disappeared. It was again displayed after movement completion as an end-point at a radial distance of 12cm. Finally, in the “veridical feedback” sub-block, the cursor remained aligned with the hand. Following the baseline block, participants underwent 280 trials of learning. Learning trials were identical to the “partial feedback” trials of the baseline sub-block, except that the cursor motion was artificially “clamped” at a fixed angle of 30° relative to the target direction (Figure 1B). Both clockwise (CW) and counterclockwise (CCW) perturbations were used with counterbalancing across participants. Participants were instructed to ignore the cursor and move their hand to the target. Cursor feedback persisted until the hand reached the midpoint (6 cm) of the distance to the target, at which point it was extinguished. This deliberate perturbation of cursor feedback, irrespective of actual hand motion, induced a consistent sensory prediction error (SPE) throughout the learning block. Upon the completion of each reach, the cursor was redisplayed at a single endpoint location at one of three predefined distances from the target: 0°, 15°, or 30°. The specific end-point angle depended on the group to which participants were assigned (n = 12 each). The distance from the target to this endpoint cursor location represented the task performance error (TPE). Importantly, this experimental design facilitated the separation of task error from sensory prediction error, where the TPE was induced by displaying end-point feedback at different locations (TPEs) while the SPE was produced through the clamped cursor feedback. Also note that the 15° and 30° TPE groups were never actually successful in making the cursor hit the target, while the 0° group was always successful (despite the SPE of 30°). Following learning, participants performed 40 washout trials, which comprised of 20 “no-feedback” and 20 “veridical feedback” trial sub-blocks, like in the baseline block. On these trials, participants were instructed to move their hand to the target.

#### Experiment 2

In experiment 2, 36 participants were recruited to one of the three groups (n =12 each) and performed reaching movements from a circular start position (0.9 cm diameter) to one of the eight targets (0.98 cm diameter) located radially at a distance of 10 cm. This task involved performing point-to-point reaches across 3 continuous blocks: baseline, learning, and washout. Cursor feedback was aligned with hand motion during the baseline and washout blocks. Both these blocks comprised of 2 sub-blocks: no feedback and veridical feedback. In the “no-feedback” sub-bock, participants performed reaches without any online cursor information, while in the subsequent “veridical feedback” sub-block, the cursor was visible and was aligned to the hand motion. Following baseline, participants performed 240 learning trials on which the target location was shifted by a predetermined angle of either 10°, 15°, or 20° (n = 12 each) once the cursor covered a 2 cm distance from the start position (Figure 2B). Target shifts were implemented by extinguishing the original target and displaying a new one at the specific location, thus creating the impression that the target had “jumped”. Target shifts could be clockwise (CW) or counterclockwise (CCW) (counterbalanced across participants). Importantly, on these trials, the cursor remained clamped in the direction of the original target, but participants were asked to ignore it and instead aim their hand to the new target. The introduction of target shifts induced TPEs. Critically, the difference in the expected motion of the cursor as they moved their hand to the new target location and its actual motion in the direction of the old target also induced an SPE. This disparity served as a catalyst for further learning induced by this newly created “online” SPE. Following learning, participants performed 40 washout trials (20 no-feedback and 20 veridical feedback). During the “no-feedback” trials, participants were instructed to avoid use of any strategies they might have used and move their hand straight to the target.

#### Experiment 3

In this experiment (n = 36), participants moved from a start position (0.9 cm diameter) to one of the eight targets (0.98 cm diameter) located at a radial distance of 10 cm. After a few familiarization trials, participants performed 40 baseline trials, where the cursor was aligned with their hand. This block was subdivided into two sub-blocks: no feedback (first 20 trials) and veridical feedback (remaining 20 trials). Following baseline, participants performed 240 learning trials. During these trials, the target was “jumped” to a new location that was either 10°, 15°, or 20° (n = 12 each) CW or CCW (counter-balanced) from the original location (Figure 3A). Participants were instructed to reach to the new target accurately. Importantly, cursor feedback remained veridical throughout the learning block. After learning, participants performed a washout block of 40 trials, which were similar to baseline (sub-blocks of 20 no-feedback trials and 20 veridical feedback trials). At the beginning of the washout block, participants were instructed to avoid using any strategy they might have developed during the learning trials and move their hand to the target directly.

**Figure 3:**
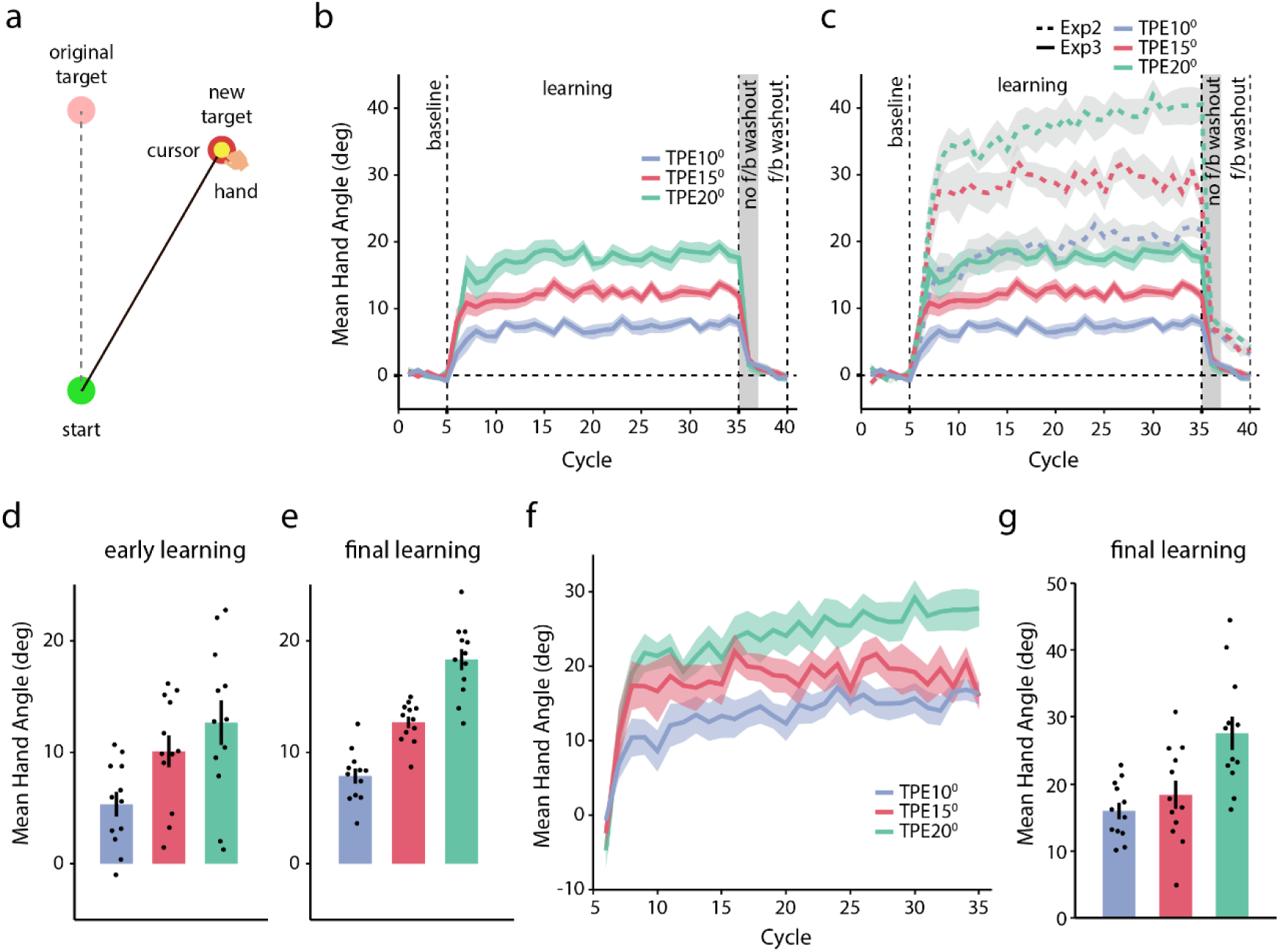
Experiment 3. **(a)** Task: As in experiment 2, the initially displayed reach target jumped by 10° (group 1), 15° (group 2), or 20° (group 3), inducing TPEs. Participants were instructed to reach to the new target. The cursor remained aligned with the hand all the time. **(b)** Group-averaged hand deviations across cycles for the three groups. **(c)** Hand deviation data from experiment 2 (dotted lines) overlaid for comparison with the data shown in (b). Shaded regions represent SEM. **(d–e)** Mean ± SEM hand deviation during (d) early learning and (e) final learning. Dots represent individual participants. **(f)** Hand deviation computed by subtracting the initial adaptation levels of Experiment 3 from all cycles of Experiment 2. **(g)** Mean hand deviation at the end of the learning period for the “learning curves” in (f).

#### Experiment 4

In experiment 4, the general experimental procedure was identical to experiment 2, except for the learning block. Participants (n = 36) were randomly assigned to three groups and performed reaching movements to one of the eight targets that appeared at a 10 cm distance. During learning trials, as participants moved a 2 cm distance, the target was “jumped” to a new location at 10°, 15°, or 20° (CW or CCW, counterbalanced) while the cursor remained clamped in the direction of the original target. While this was a general feature of the entire learning block, the learning trials appeared in sub-blocks that differed in terms of what the participants were expected to do. These sub-blocks were presented in an alternating set of 16 trials each (Figure 4A). In the first “learning” sub-block, participants were asked to ignore the cursor and aim their hand to the new target location. As in experiment 2, this was expected to lead to a rapid change in hand angle as participants deliberately aimed away, which would then additionally induce an SPE and drive further changes in hand angle. To probe whether this was indeed occurring from the very outset of the learning block, and to understand the broader time-course of SPE-mediated learning, we included a second “test” sub-block, during which participants were asked to ignore the target shift and aim to the original target location. Moreover, on these “test” trials the cursor feedback was eliminated. This enabled us to understand *when* the changes in hand angle mediated by the SPE induced online on the “learning” sub-block trials were coming into effect. Following the learning block, participants performed 40 washout trials similar to those in experiments 2. During these trials, they were instructed to move their hand directly to target avoiding any aiming strategy that they might be using during learning.

**Figure 4:**
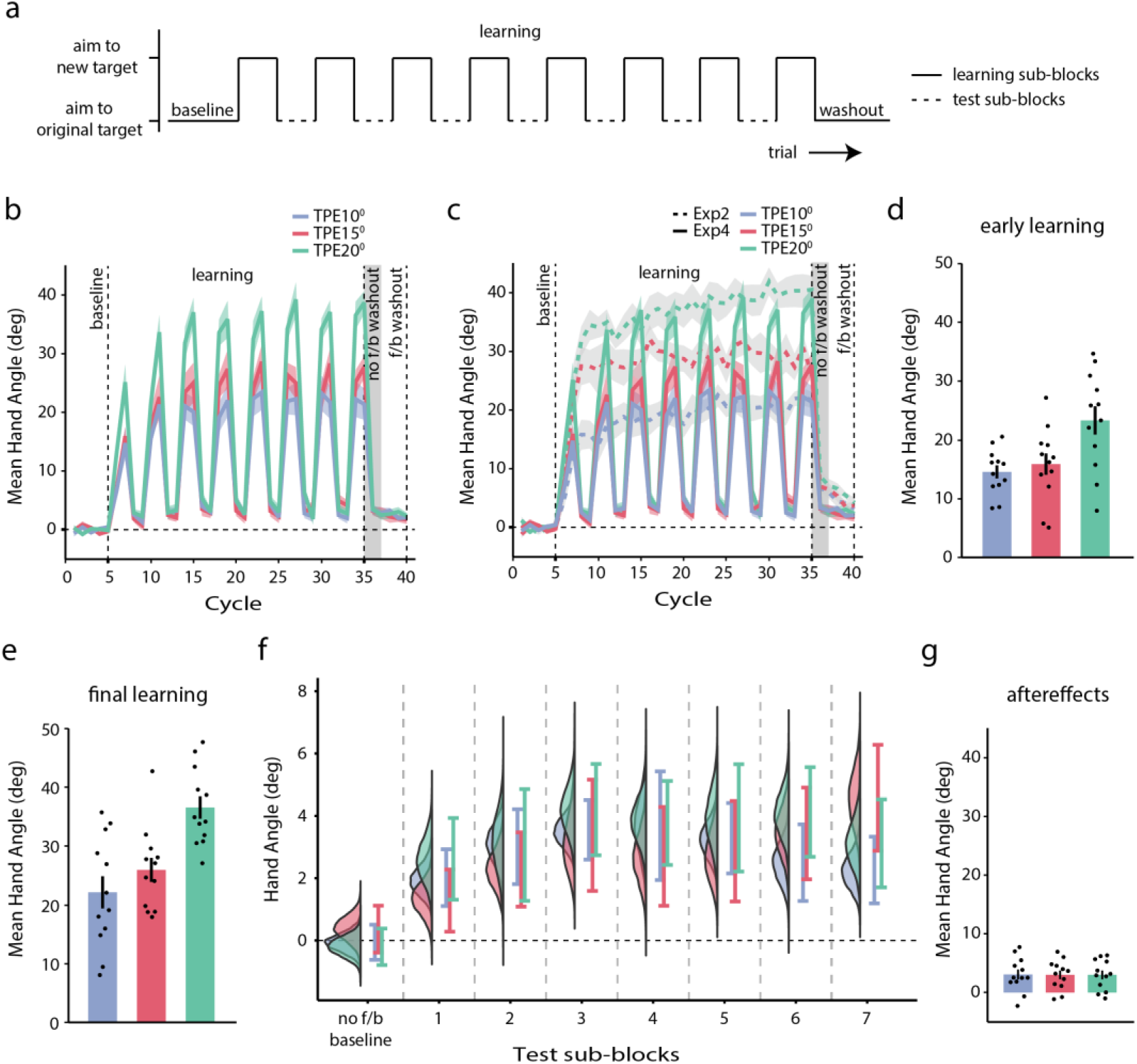
Experiment 4. **(a)** Task: Participants performed baseline (no perturbation movements) followed by a learning block composed of alternating sub-blocks of learning and test trials. The learning sub-blocks were identical to those Experiment 2, wherein participants reached to a “jumped” target while cursor feedback remained clamped in the direction of the original target. During the test sub-blocks, cursor feedback was withheld, and participants were instructed to aim at the original target location. Learning and test sub-blocks thus differed in terms of whether participants aimed for the new target location or the original one. **(b)** Group-averaged hand deviations (calculated relative to the original target direction) across cycles for the three groups. **(c)** Hand deviation data from experiment 2 (dotted lines) overlaid for comparison with the data shown in (b). Shaded regions represent SEM. **(d–e)** Mean ± SEM hand deviation during (d) early learning, and (e) final learning. **(g)** Distributions of bootstrapped means and 95% confidence intervals of the hand deviation for each group across the seven no-feedback test sub-blocks. **(f)** Mean ± SEM hand deviation during no-feedback washout. Dots represent individual participants.

### Data analysis

Recorded X-Y hand position data were first filtered using a Butterworth filter with a cutoff frequency of 10 Hz. Velocity values were subsequently obtained by differentiating the filtered position data. The primary dependent variable in this study was the change in hand angle across trials. Hand angle was defined as the angle between two vectors: one connecting the start position to the target, and the other connecting the start position to the hand position at peak velocity. To account for individual baseline biases, the mean baseline hand angle for each participant was subtracted from the hand angle values of that participant from all trials. Motor adaptation was quantified as change in these baseline-corrected hand angles across the course of learning trials. Participants who failed to learn and trial where the movement was not initiated excluded from further analysis. Further, hand angle values beyond the range ±100° were also excluded. Early learning was defined as the mean direction error over the first 32 learning trials, i.e., 8 cycles in experiment 1 and 4 cycles in experiment 2, 3 and 4. Similarly, final learning was also defined as mean direction error over the last 32 learning trials. Aftereffects magnitude was quantified as average of the first 16 trials of the no-feedback washout sub-block. In experiment 1, we also calculated learning rate by first bootstrapping the learning data and then fitting an exponential of the form [y = ae^-bx^ + c] to each bootstrapped sample. For the statistical comparisons one-way ANOVA, one sample t-tests and linear mixed-effects model were used. For the linear mixed-effects model, the test sub-blocks and group were used as fixed-effects, and participants were used as random-effects, allowing them to have random intercepts and slopes. The significance threshold was set at 0.05.

## RESULTS

### Experiment 1

In our first experiment, 3 groups of participants participated in a modified error-clamp paradigm in which we extinguished the clamped feedback cursor at the halfway point to the target (Figure 1B). After the full target distance had been covered, we re-displayed the cursor at either 0° (TPE0° group), 15° (TPE15° group) or 30° (TPE30° group) from the target. Thus, while the SPE remained fixed, the TPE could be varied. We compared the change in hand angle (hand deviation) and learning rate across the three groups to probe whether the TPEs of different magnitudes would influence the amount of learning driven by the SPE. Broadly, we found no group differences in adaptation.

Figure 1C shows that all three groups displayed typical adaptation curves and reached asymptote with repeated practice. There was no difference (F_2,33_ = 0.29, p = 0.75, Ƞ^2^_p_ = 0.017) in the early change in hand angle across groups (Figure 1D, mean ± sem: TPE0° = 7.023 ± 0.560, TPE15° = 6.952 ± 1.178, TPE30° = 6.142 ± 0.880). Likewise, the final level of learning (Figure 1E) was comparable across the three groups (mean ± sem: TPE0° = 20.345 ± 1.455, TPE15° = 20.166 ± 1.743, TPE30° = 18.995 ± 1.852; F_2,33_ = 0.188, p = 0.83, Ƞ^2^_p_ = 0.011). This preliminarily indicated that the magnitude of the TPE did not influence learning in our task. To probe this better, we also compared aftereffects across the groups. Again, we found no group differences (Figure 1F, F_2,33_ = 0.166, p = 0.848, Ƞ^2^_p_ = 0.01), indicating that adaptation remained invariant despite variations in the TPE. Could TPEs have influenced the rate of learning than just the initial and final hand angle or aftereffects? To address this, we compared the learning rate across the three groups. We fit a single rate exponential function of the form [y = ae^-bx^ + c] to 10000 bootstrapped samples of hand deviation for each group and obtained the learning rate. However, there were no group differences as indicated by the overlap in the 95% confidence intervals of the bootstrapped distributions (TPE0° = 0.0672– 0.1181, TPE15° = 0.0503–0.1100, TPE30° = 0.0676–0.1152). Collectively, the findings of this experiment suggested that TPEs of different magnitudes did not differentially influence adaptation to the SPEs induced by the error-clamp.

### Experiments 2 and 3

In our second experiment, TPEs were induced by jumping the reach target by 10° (TPE10°), 15° (TPE15°) or 20° (TPE20°) following movement initiation, but the cursor was clamped in the direction of the original target (Oza et al., 2024). This design produced a TPE as the cursor always missed the jumped target location (Figure 2B). Participants were asked to respond by aiming their hand to the new target location, but since the cursor was clamped towards the original target, an SPE was also induced. Note that this SPE came about due to the mismatch between the expected and actual direction of cursor motion; that is, participants expected the cursor to go with the hand towards the new target, but it actually traveled in the direction of the original target. Our main question of interest was whether learning driven by SPEs would differ across our three groups.

Figure 2C shows the learning curves for the 3 groups. Participants in all groups demonstrated rapid changes in hand deviation in the direction of the new target. Remarkably, within a few trials, hand deviation exceeded the angle by which the target had been jumped; this trend to “overcompensate” beyond the jump magnitude was evident across all 3 groups (Figure 2D, mean ± sem: TPE10° = 12.151 ± 1.699, TPE15° = 20.884 ± 2.640, TPE20° = 24.462 ± 2.252), and group differences at the early learning phase were statistically significant (F_2,33_ = 8.059, p < 0.001, Ƞ^2^_p_ = 0.328). As participants continued to aim for the new target, the induced SPE caused further changes in hand angle, albeit more gradually in all groups. By the end of learning, all participants demonstrated significant hand deviation with notable differences evident between groups (Figure 2E, mean ± sem: TPE10° = 21.314 ± 1.223, TPE15° = 28.468 ± 2.091, TPE20° = 40.245 ± 2.490; F_2,33_ = 22.71, p < 0.001, Ƞ^2^_p_ = 0.579). We also observed a non-zero after-effect during the early washout trials (Figure 2F), likely arising from implicit learning triggered by this SPE (Oza et al., 2024). Notably, there were no group differences in its magnitude (mean ± sem: TPE10° = 6.394 ± 0.894, TPE15° = 6.752 ± 0.900, TPE20° = 7.927 ± 0.510; F_2,33_ = 0.649, p = 0.529, Ƞ^2^_p_ = 0.038), suggesting again that there was no impact of TPE size on SPE-based learning.

However, we wondered whether there might be subtle differences in the SPE-mediated learning that might be sensitive to TPE magnitude that the after-effect might not have captured. If so, then group differences would be evident when the SPE-mediated learning is isolated from the combined effect of the requirement to aim to the new target and the subsequent adaptation to the SPE induced thereby. To begin isolating the SPE-mediated component, in our third experiment, we first measured the change in hand angle that would result from re-aiming to a new target location. To do so, we measured the change in hand angle in response to target jumps alone under conditions of veridical (instead of clamped) cursor feedback (Figure 3A).

Three groups of participants performed point-to-point reaching movements with the cursor aligned accurately with actual hand motion. During learning trials, the target was shifted by 10° (TPE10°), 15° (TPE15°), or 20° (TPE20°) to induce TPEs, and participants were instructed to reach to the new target location. Across all groups, participants rapidly adjusted their hand trajectories toward the shifted target (Figure 3B). Hand deviation was evidently different across the groups during the early learning phase itself (Figure 3D, mean ± sem: TPE10° = 5.346 ± 1.103, TPE15° = 10.094 ± 1.423, TPE20° = 12.654 ± 1.996; F_2,33_ = 5.708, p = 0.007, Ƞ^2^_p_ = 0.257). Participants showed near complete compensation of the target jump within a few trials, and only incremental changes in hand angle thereafter. At the end of learning (Figure 3E), the hand deviation reached 7.872 ± 0.672, 12.692 ± 0.521, 18.305 ± 0.941 on average for the 10°, 15° and 20° groups respectively. The difference in hand angle across the three groups at the end of learning was statistically reliable (F_2,33_ = 50.833, p < 0.001, Ƞ^2^_p_ = 0.755). This indicated that participants effectively adapted their movements to account for the jump-induced TPE (Sadaphal et al., 2022). Importantly, unlike experiment 2, no overcompensation was observed in response to target shifts.

Since our larger aim was to isolate the SPE-mediated learning in experiment 2, we proceeded to subtract out, for each jump magnitude separately, the average TPE-mediated learning of the early learning phase of experiment 3 from the learning observed in experiment 2. We reasoned that this would be akin to removing, from the learning data of experiment 2, the component that was solely due to the explicit re-aiming to the new target location. Upon doing so, we obtained “learning curves” in which group differences were still evident (Figure 3F), and also statistically significant at the end of the learning phase (F_2,33_ = 9.354, p < 0.001, Ƞ^2^_p_ = 0.362, Figure 3G).

It could be argued that this reflects differences in SPE-mediated learning, as a function of TPE magnitude. In other words, these group differences might provide evidence for the modulation of implicit learning by task errors. However, we also noted that the removal of the jump-mediated component did not zero out the overcompensation observed in the early learning phase of experiment 2. On average, there was an “additional” hand deviation of 6.805°, 10.790° and 11.808° for the 10°, 15° and 20° target jumps respectively that was still present during the early stages of experiment 2. Moreover, it was unclear whether this overcompensation resulted from continued engagement of deliberative processes in response to the jump-induced TPE (participants trying to “pull” the clamped cursor towards the new target location) or from the SPE induced by the mismatch between hand and cursor motion. Recall that our goal was to remove the TPE-mediated component completely, isolate the SPE-mediated component, and investigate whether this SPE-driven learning was influenced by the TPE. Clearly, this subtractive approach was limited in doing so. It therefore became imperative to find a different approach to isolate implicit learning even if participants were overcompensating initially. We proceeded to do so in experiment 4.

### Experiment 4

The experimental design for our fourth and final experiment was similar to that of experiment 2 but included intermittent sub-blocks of 16 test trials each alternating with learning sub-blocks of 16 trials each (Figure 4A). On the test trials, the cursor feedback was eliminated, and participants were instructed to ignore the target jump and reach to the original target location. Inserting these trials was important since it enabled us to probe the evolution of SPE-mediated changes in hand angle – we expected that as adaptation progressed, participants would show a bias towards the trained direction (rather than moving in the direction of the original target) on these test trials. At the same time, the learning sub-blocks allowed us to obtain overcompensation identical to experiment 2 in the early learning phase, but how this overcompensation came about would be irrelevant. Rather, our key comparison would be the magnitude of the bias across the different groups.

Figure 4B shows the change in hand angle for the three groups during the learning and test sub-blocks. As is evident, there was rapid and robust overcompensation in all groups, with the average change in hand angle being 14.569 ± 1.113°, 15.887 ± 1.815°, 23.337 ± 2.405° during the early learning phase for the 10° (TPE10°), 15° (TPE15°), and 20° (TPE20°) jump groups respectively (Figure 4D). As learning progressed, there was an increasing trend in the hand deviation. At the end of learning, hand angle had increased to 22.155 ± 2.730°, 25.956 ± 2.040° and 36.574 ± 1.894° for the TPE10°, TPE15°, and TPE20° groups respectively (Figure 4E). These results were all consistent with the general trend seen in experiment 2.

We then turned our attention to the test sub-blocks. Firstly, we obtained a small but non-zero “after-effect” on the first test sub-block (1.956 ± 0.488°, 1.311 ± 0.525°, 2.590 ± 0.708° for the TPE10°, TPE15°, and TPE20° groups respectively (Figure 4F). This after-effect was significantly different from zero in all cases (TPE10°: t(11) = 4.006, p = 0.002, Cohen’s d = 1.156; TPE15°: t(11) = 2.497, p = 0.03, Cohen’s d = 0.721; TPE20°: t(11) = 3.658, p = 0.004, Cohen’s d = 1.056), suggesting that changes in hand angle driven by the SPE had begun to occur during the early learning phase itself. To reiterate, this SPE was induced by the participants’ expectation that the cursor would go with the hand as they aimed to the new target location, while its actual motion remained clamped in the direction of the original target on the learning trials. Importantly, there were no group differences in hand deviation on the first test sub-block (F_2,33_ = 1.208, p = 0.312, Ƞ^2^_p_ = 0.068). This suggested that the SPE-mediated effects were not affected by the TPE magnitude during early learning.

Figure 4F shows the magnitude of the hand deviation for each group for each of the test sub-blocks, reflecting the time course of implicit learning development with training. As can be seen, there was an increasing trend in hand deviation over the first 3 or so sub-blocks for all groups, which was also confirmed using a linear mixed-effects model (fixed effect slope = 0.51, [0.22,0.80], p = 0.001, Cohen’s f^2^ = 0.06). However, there was no significant effect of group or interaction between block and group (p>0.05). The magnitude of hand deviation stabilized after this initial phase for all groups. This mirrored the canonical, SPE-driven change in hand angle that tends to gradually increase over the initial set of learning trials and asymptotes thereafter.

We next asked whether there were any group differences in hand deviation at each sub-block. We did not find this to be the case. Table 1 shows the 95% confidence intervals for hand deviation for each group for each of the 7 test sub-blocks (in addition to the no-feedback baseline sub-block), derived from 10000 bootstrapped samples of the data. The overlap in confidence intervals of the groups at each time point suggests that the degree of implicit adaptation was not different between them even though they experienced different TPEs. In other words, the amount of implicit learning was unaffected by TPE magnitude. Finally, we compared the post-learning after-effect (Figure 4G) and again found no group differences (F_2,33_ = 0.002, p = 0.998, Ƞ^2^_p_ = 1.361 x 10^-4^), pointing to the fact that implicit learning did not vary with the size of the TPE.

**Table 1:** 95% confidence interval values of 10000 bootstrapped samples for the three groups at each test sub-blocks.

| Group | no f/b<br>baseline | sub-block 1 | sub-block 2 | sub-block 3 | sub-block 4 | sub-block 5 | sub-block 6 | sub-block 7 |
| --- | --- | --- | --- | --- | --- | --- | --- | --- |
| TPE10° | -0.6229,<br>0.5037 | 1.1058,<br>2.9293 | 1.8079,<br>4.2131 | 2.5975,<br>4.5099 | 1.9379,<br>5.4247 | 2.1515,<br>4.4116 | 1.2669,<br>3.7276 | 1.1873,<br>3.3311 |
| TPE15° | -0.3903,<br>1.1144 | 0.2801,<br>2.2760 | 1.0847,<br>3.4704 | 1.5882,<br>5.1579 | 1.1141,<br>4.2850 | 1.2502,<br>4.4778 | 1.9634,<br>4.9070 | 2.8824,<br>6.2805 |
| TPE20° | -0.7928,<br>0.3811 | 1.3069,<br>3.9328 | 1.2699,<br>4.8573 | 2.7374,<br>5.6636 | 2.4242,<br>5.1159 | 2.2134,<br>5.6547 | 2.6865,<br>5.5590 | 1.7063,<br>4.5287 |

## DISCUSSION

For the sensorimotor system to operate effectively, it must learn from errors. During movement, such errors arise from at least two sources: a prediction about the sensory consequences of the action that differs from the actual feedback received (SPE), and a failure to achieve the desired task goal (TPE). In this work, we focused on the interaction between these error sources during learning. For this purpose, we employed two new error-clamp tasks, each focusing on different shortcomings of the traditional error-clamp learning paradigm. Our experimental findings suggest that SPE-mediated learning remains robust and unaffected by TPE manipulations, thereby pointing to a certain imperviousness of implicit motor learning to task outcomes.

An open issue in motor adaptation research is whether the typical error-clamp paradigm truly isolates the SPE-mediated changes in motor plans, or inadvertently allows task performance (TPE) to influence behaviour. This issue arises because it is not possible to assess whether a participant has indeed completely ignored the TPE despite explicit instructions to do so. Early variability in hand movement direction, such as abrupt shifts in movement direction opposite to the clamped cursor, suggests that participants might not be able to fully suppress TPE processing. Such adjustments align with the known capacity of a TPE to drive strategic corrections (McDougle et al., 2015; McDougle & Taylor, 2019; Morehead et al., 2015; Sadaphal et al., 2022). Critically, the persistence of such behavior in trials preceding significant SPE-driven adaptation challenges the assumption that clamped paradigms fully dissociate these error signals. Our study circumvented this limitation through two novel approaches: in experiment 1 we assumed that the TPE remains salient in standard clamps and manipulated the TPE magnitude independent of SPE. In experiments 2 and 4 we bypassed the inherent TPE of error-clamps by inducing TPEs via target jumps, while also creating an SPE on the fly. Thus, by designing tasks that enabled us to independently test both scenarios, i.e., TPE as a contributor versus an ignored signal, we are now able to provide a dual-lens framework to resolve this issue. The robustness of SPE-mediated learning across both paradigms, despite divergent TPE manipulations, strongly supports its primacy in implicit adaptation without any interaction with the TPE. This of course does not mean that the TPE carries no influence. As a number of recent results, including some of our own have shown, the TPE might set in motion a completely distinct mechanism - volitional, explicitly-accessible adjustments to reach direction (McDougle et al., 2015; McDougle & Taylor, 2019; Morehead et al., 2015; Sadaphal et al., 2022).

Our findings appear to contradict some recent results showing that trial-by-trial learning rates, measured as hand deviation changes between consecutive trials, are modulated by TPE magnitude (Al-Fawakhiri et al., 2023; Tsay et al., 2022). This apparent contradiction brings forth a critical nuance, that the temporal structure of errors (blocked vs. random) may determine whether the TPE influences SPE-mediated learning. In our block design, with SPE and TPE relationships fixed across trials, the motor system may treat errors as stable contexts. Under such conditions, SPE-driven adaptation aligns with the implicit mechanism described in classical studies (Mazzoni & Krakauer, 2006; Morehead et al., 2017; Shadmehr & Mussa-Ivaldi, 1994), prioritizing improved sensory prediction over task success. In contrast, in trial-by-trial designs where error magnitudes and their relationships change continuously, TPEs may be interpreted as a dynamic contextual signal that could influence SPE-driven learning. The COIN model (Heald et al., 2021) might be useful in formalizing this idea: when context transition probabilities are high (e.g., random perturbations), the motor system weights TPE more heavily to rapidly adjust strategies, modulating single-trial learning. This could explain why Tsay and colleagues (Tsay et al., 2022) observed TPE-driven modulation, whereas our block design, with low context transition probabilities, revealed no such interaction. In sum, this implies that in stable environments, SPE-driven recalibration dominates, rendering learning indifferent to TPE magnitude. In contrast, in rapidly changing contexts, the TPE might gate explicit adjustments without altering SPE-based adaptation. Our results thus refine, rather than contradict, prior work, highlighting that the role of the TPE is likely to be context-dependent and largely orthogonal to SPE-driven learning.

Our paradigms also address some inconsistencies between theoretical framing and computational models of the role of TPEs in error clamp paradigms. Specifically, theoretical models suggest that error-clamp paradigms isolate SPE by eliminating or minimizing effects of the TPE (Morehead et al., 2017). That is, the TPE plays no role in learning under these conditions. However, corresponding computational models include a TPE term (Kim et al., 2019; Tsay et al., 2022). For example, in Experiment 4 of Tsay et al. (2022), the target was jumped to different locations while the cursor remained clamped in a fixed direction away from the target. When modeling the ensuing behavior, the TPE was included as the distance between the clamped cursor and the shifted target. Moreover, this TPE included contributions from both the clamp and the target jumps (at least in their “jump away” conditions), which adds an additional layer of complexity to the interpretation, as the TPE induced by the rotation (i.e., the clamp) and the TPE induced by a goal shift (i.e., target jump) could engage distinct cognitive or neural mechanisms (Diedrichsen et al., 2005; Mutha et al., 2011a, 2011b). Given this, combining TPEs from multiple origins into a single term within a model, risks obscuring the specific contributions of each type. In our study, this issue was minimized by the use of two distinct task designs: one involving perturbation-linked TPEs, and another involving jump-induced TPEs. As stated earlier, we found no evidence that either type of TPE modulates SPE-driven implicit learning.

Experiments 2 and 4 demonstrated systematic overcompensation in TPE-mediated re-aiming behavior. When target jumps were used to introduce TPEs and cursor feedback remained clamped to the original target location, participants consistently adjusted their hand movements beyond what was needed to hit the target. In fact, even after accounting for the SPE-mediated learning, the residual overcompensation magnitudes substantially exceeded the actual TPE magnitudes. This effect likely stemmed from the lack of success signals (cursor hitting the target) as participants re-aim. Indeed, in Experiment 3, where true cursor feedback was provided, participants did not demonstrate any overcompensation, which was also the case in our past work (Sadaphal et al., 2022). Without this feedback, participants perhaps kept refining their aim, pausing only when proprioceptive feedback may have signaled excessive hand deviation relative to the jump magnitude. Implicit recalibration was likely simultaneously triggered, driven by the SPE. Importantly, the prior overcompensation did not influence the amount of SPE-driven adaptation. We found that the despite differences in the magnitude of TPE-mediated compensation, the after-effects remained indistinguishable in experiments 2 and 4. This again provided confirmation of the idea that that SPE-driven recalibration remains fundamentally independent of strategic adjustments driven by target jump mediated TPEs.

More broadly, the dissociation between TPEs and SPEs reflects two complementary processes that contribute to motor learning. TPEs, which signal deviations from task goals leading to activation in reward-sensitive cortico-striatal pathways (Diedrichsen et al., 2005; Inoue et al., 2016), engage explicit strategy adjustments akin to model-free processes involving trial-and-error refinement of actions to achieve immediate success (e.g., re-aiming or mental rotation). Such adjustments prioritize direct behavioral outcomes over internal representations of the environment. In contrast, SPEs, arising from discrepancies between predicted and actual sensory feedback, involve processing within cerebellum (Galea et al., 2011; Martin et al., 1996; Morehead et al., 2017) and posterior parietal cortex (Kumar et al., 2020; Mutha et al., 2011b, 2017), and drive updates to internal models of limb-environment interactions, aligning with model-based processes. Here, adjustments are guided by predictive accuracy rather than task success, enabling long-term improvements in sensorimotor control. While the model-free system rapidly adapts actions to meet task demands, the model-based system incrementally refines predictive models to optimize future movements. These processes likely operate in parallel: TPE-driven strategies adjust actions to minimize task failure, while SPE-driven adaptation ensures movements remain calibrated to the body’s biomechanics and environmental constraints. Collectively, these mechanisms enable robust motor learning across diverse contexts, from immediate goal achievement to sustained environmental interaction.

## Acknowledgements

We thank Adarsh Kumar for many helpful discussions regarding this work. This work was supported by grants from the Department of Science and Technology, Government of India. Support from IIT Gandhinagar is also acknowledged.

